# A Flow Procedure for the Linearization of Genome Sequence Graphs

**DOI:** 10.1101/101501

**Authors:** David Haussler, Maciej Smuga-Otto, Benedict Paten, Adam M Novak, Sergei Nikitin, Maria Zueva, Dmitrii Miagkov

## Abstract

Efforts to incorporate human genetic variation into the reference human genome have converged on the idea of a graph representation of genetic variation within a species, a genome sequence graph. A sequence graph represents a set of individual haploid reference genomes as paths in a single graph. When that set of reference genomes is sufficiently diverse, the sequence graph implicitly contains all frequent human genetic variations, including translocations, inversions, deletions, and insertions.

In representing a set of genomes as a sequence graph one encounters certain challenges. One of the most important is the problem of graph linearization, essential both for efficiency of storage and access, as well as for natural graph visualization and compatibility with other tools. The goal of graph linearization is to order nodes of the graph in such a way that operations such as access, traversal and visualization are as efficient and effective as possible.

A new algorithm for the linearization of sequence graphs, called the flow procedure, is proposed in this paper. Comparative experimental evaluation of the flow procedure against other algorithms shows that it outperforms its rivals in the metrics most relevant to sequence graphs.

## 2 Motivation

The current human reference genome consists essentially of a single representative of each of the human chromosomes. In essence, an arbitrary person’s genome is chosen to represent all of humanity. This leads to loss of information and bias. Efforts to incorporate human genetic variation into the reference human genome have converged on the idea of a graph representation of genetic variation within a species, a genome sequence graph [1].

In its mathematically most simple form, each node of a sequence graph contains a single DNA base that occurs at an orthologous locus in one or more of the haploid genomes represented in the graph. Each arc represents an adjacency (chemically, a covalent bond) that occurs between consecutive instances of bases in those genomes. Excluding reversing joins (see below) each arc is directed according to the default strand direction of the DNA sequence used to build the graph, connecting the 3’ side of the previous base (the tail of the edge) with the 5’ side of the next base (the head of the edge). At points where the individual genomes differ to the right (i.e., 3’, downstream) of an orthologous base, the node representing that base will have two or more outgoing arcs. For example, the graph in Figure 2 was built from the DNA strands *CATCGCT*, *CATCGT, CAGCGAT* and *CATCGAGAGAGCT* aligned at orthologous bases as shown in Figure 1.

**Figure 1.**
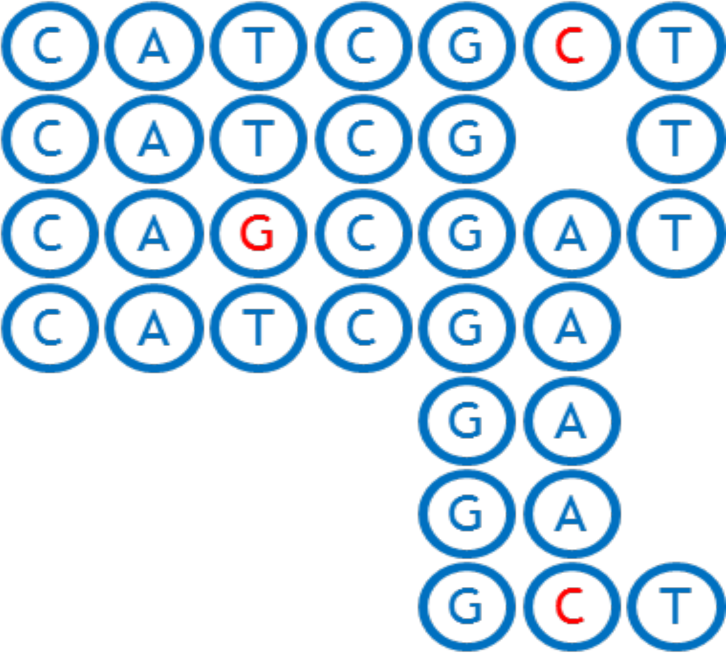
Alignment of CATCGCT, CAGCGAT, CATCGAGAGAGCT DNA strands.

**Figure 2.**
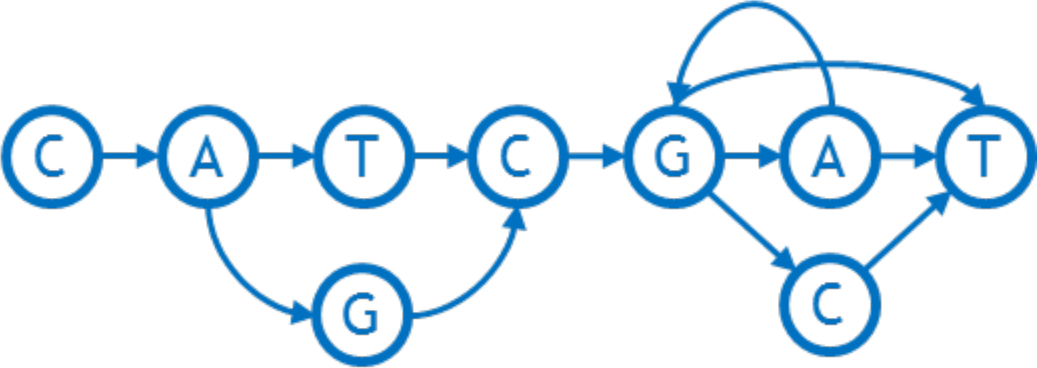
An example of a sequence graph reflecting SNP (T/G), tandem duplication GA -> GAGAGA, and deletion/SNP (A/C/-).

Representing a genome as a graph requires building a number of tools to work with it efficiently. In particular, one needs to linearize the graph, that is order the nodes from left to right in a straight line. Linearization facilitates visual perception of a graph, allows software to index the nodes in a familiar manner, and imposes natural order of bases useful in storage, search and analysis, for example enabling traversal from left to right with minimum feedback runs. Figures 3 and 4 show examples of a linearized graph and individual genomes in it. Here we have also added a new feature to the graphs in the form of **arc weights**. An arc weight is used to signify the importance of an arc in typical applications running on the graph. Normally we set the arc weight to the number of times that the arc is traversed in the reference genomes used to build the graph, under the assumption that arcs used frequently in the reference genomes 3 will also be used frequently in the applications of the sequence graph built from them. Some reference genomes may be weighted more than others.

**Figure 3.**
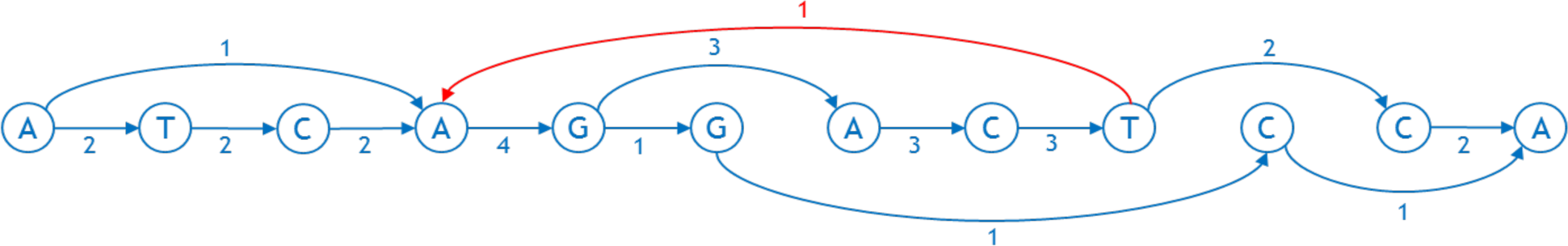
An example of a linearized sequence graph with weighted arcs. The arc in red is directed from the right to the left and is called feedback arc.

**Figure 4.**
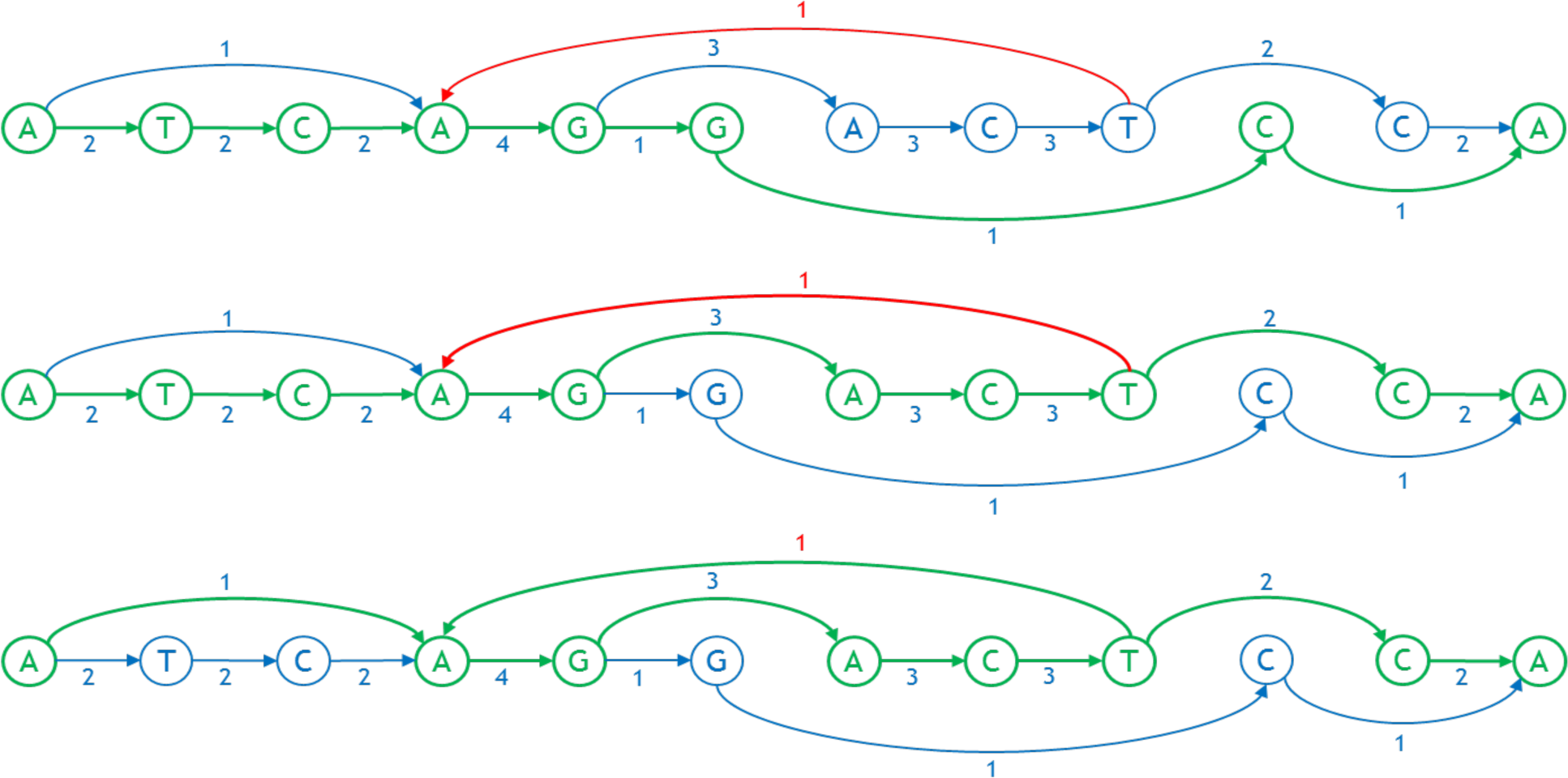
Three individual genomes as paths in the linearized graph, shown in green. The first and second genomes are ATCAGGCA and ATCAGACTCA, respectively. The third one is AAGACTAGACTCA where the arc between T and A is a feedback arc.

## 3 Problem Statement

A linearization of a sequence graph aims to make the total weight of all feedback arcs, called the **weighted feedback** (see Figure 3), small, along with the “width” (number of arcs) crossing any vertical line in the layout (called a “cut”, see Figure 5). Unnecessary feedback arcs make many types of genetic analysis more inefficient, as these typically proceed left-to-right on a conventional reference genome. An arc crossing a cut is considered to be part of an allele spanning that cut (see [2]), so a graph with smaller cut width at a cut has fewer alleles at that cut. The mean of the cut width over all cuts in the graph is called the **average cut width**.

**Figure 5.**
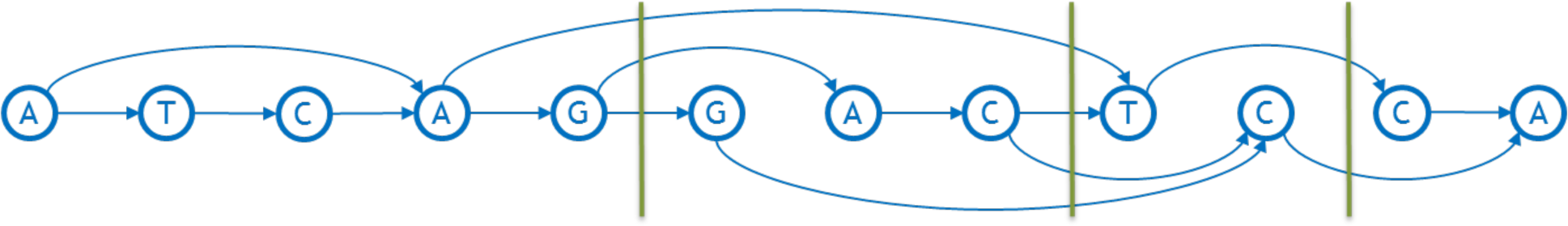
Examples of cuts and cut widths. Three cuts are shown with green vertical lines. Their widths are, from left to right, 3, 4, and 2 respectively.

In light of their importance for genetic analysis, we will evaluate a linearization based on its average cut width and either the weighted feedback or the number of feedback arcs it contains.

Unfortunately, the problems of minimizing weighted feedback, and of minimizing the average cut width, are each separately difficult. The simpler problem of minimizing the number of feedback arcs is known in literature as the *feedback arc set problem*, or FAS for short [3]. This is an NP-hard problem [4], but there are a number of various heuristic approaches to approximating a solution to it [5]. The problem of minimizing the average cut width [6] is also known to be NP-hard. A good heuristic [7] is necessary for good results. In our case, starting our procedure with the “primary path” taken by the reference genome is a natural choice. This is the first time to our knowledge that a heuristic algorithm to minimize both metrics at the same time has been proposed.

## 4 Algorithm Description

We propose here a simple heuristic divide-and-conquer approach to linearly order the bases of a graph that tries to achieve either small weighted feedback (or small number of feedback arcs) and small average cut width. The key algorithmic tool is max-flow/min-cut in a directed graph [8], so we call it the **flow procedure**. Prior to applying the flow procedure, the sequence graph is “groomed” as described at the end of this section.

### 4.1 First Step. Find the Backbone

Following grooming, the flow procedure starts with a connected graph with directed arcs and a designated linear ordering of a subset of the bases called the **backbone**. Arcs leading from the backbone to nodes not on the backbone are called **out-arcs**, and arcs directed into the backbone from nodes not on the backbone are called **in-arcs**. Grooming guarantees such the first base of the backbone has no in-arcs and the last has no out-arcs. Extra dummy bases are added at either end if necessary. Normally the initial backbone is a biologically determined primary path of the graph, e.g. from a selected haploid reference genome in the set of genomes used to build the graph, perhaps the existing haploid reference human genome. The flow procedure creates a backbone using its internal heuristics if none is given *a priori.* In linearizing the sequence graph by creating a total ordering of the bases, the relative order of the bases in backbone will not change. The rest of the bases in the graph will be inserted either between bases of the backbone, before the backbone, or after the backbone. Thus, any feedback arcs already in the backbone ordering, called **initial feedback arcs**, will remain as feedback arcs in the final ordering.

Consider the graph depictured in Figure 6. The backbone is CGATC horizontally across the middle (highlighted by dark blue color). The three out-arcs of the backbone are shown with thick green arrows. The two in-arcs are shown with thick purple arrows. The weights are assumed to reflect usage, but we also assume the usage statistics may be partial. Hence the weight coming into a base does not always match the weight coming out.

**Figure 6.**
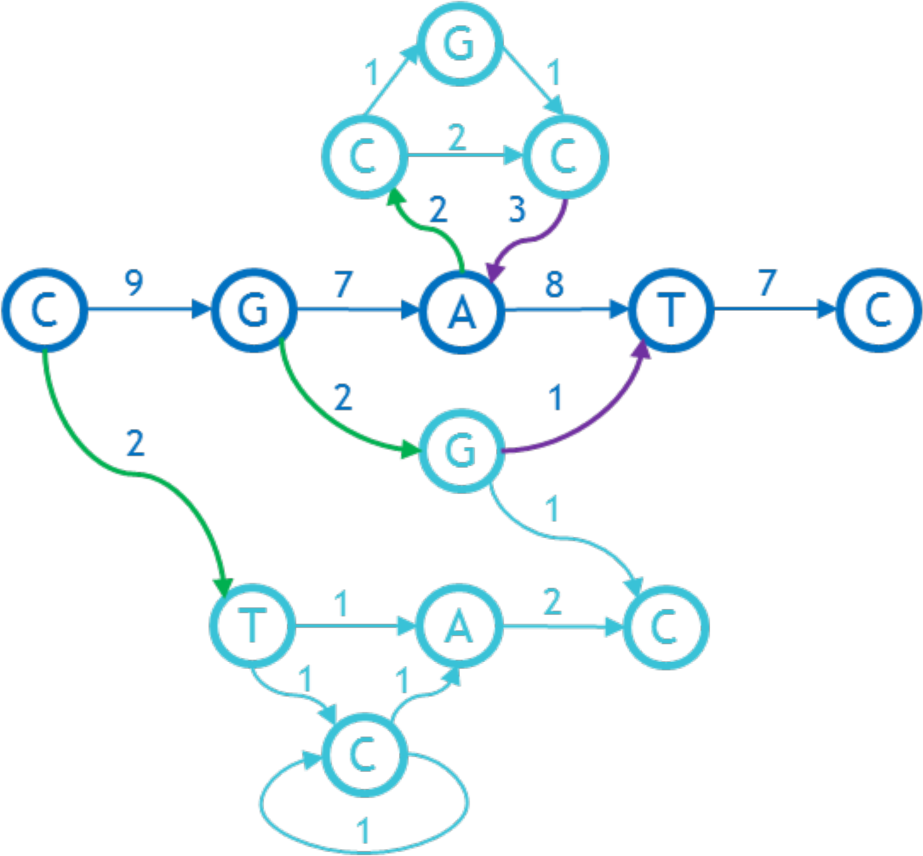
An initial directed graph with weights on the arcs.

### 4.2 Second Step. Add Source and Sink

We set up a max flow/min cut network as follows. The nodes and arcs of the network are nodes and arcs of the graph. The capacity of the arc is its weight. In addition, there is a special source node and a special sink node. The network arcs for these are defined as follows: we let *N* be the maximum of the sum of the weights of the outgoing arcs for any node in the graph and we add an arc of capacity *N* + 1 from the source node to each base on the backbone that has an out-arc. Then each in-arc on the backbone is redirected to the sink node with no change in capacity.

### 4.3 Third Step. Determine the Minimum Cut and Delete It

The maximum flow in our example is easily seen to be 3, and it maximizes the capacity of the two arcs that are cut by the green bars (Fig. 7). Therefore, these form a minimum cut. Since the capacities of the arcs connecting the source to the backbone are too high to be achieved by any flow, none of these are in the cut. Therefore, the cut must split the flow network into an **out-component** containing the source and its outgoing arcs, and an **in-component** containing the sink. In Figure 7 the in-component consists of the uppermost nodes *C, G, C* and the sink, and the out-component is the remainder.

**Figure 7.**
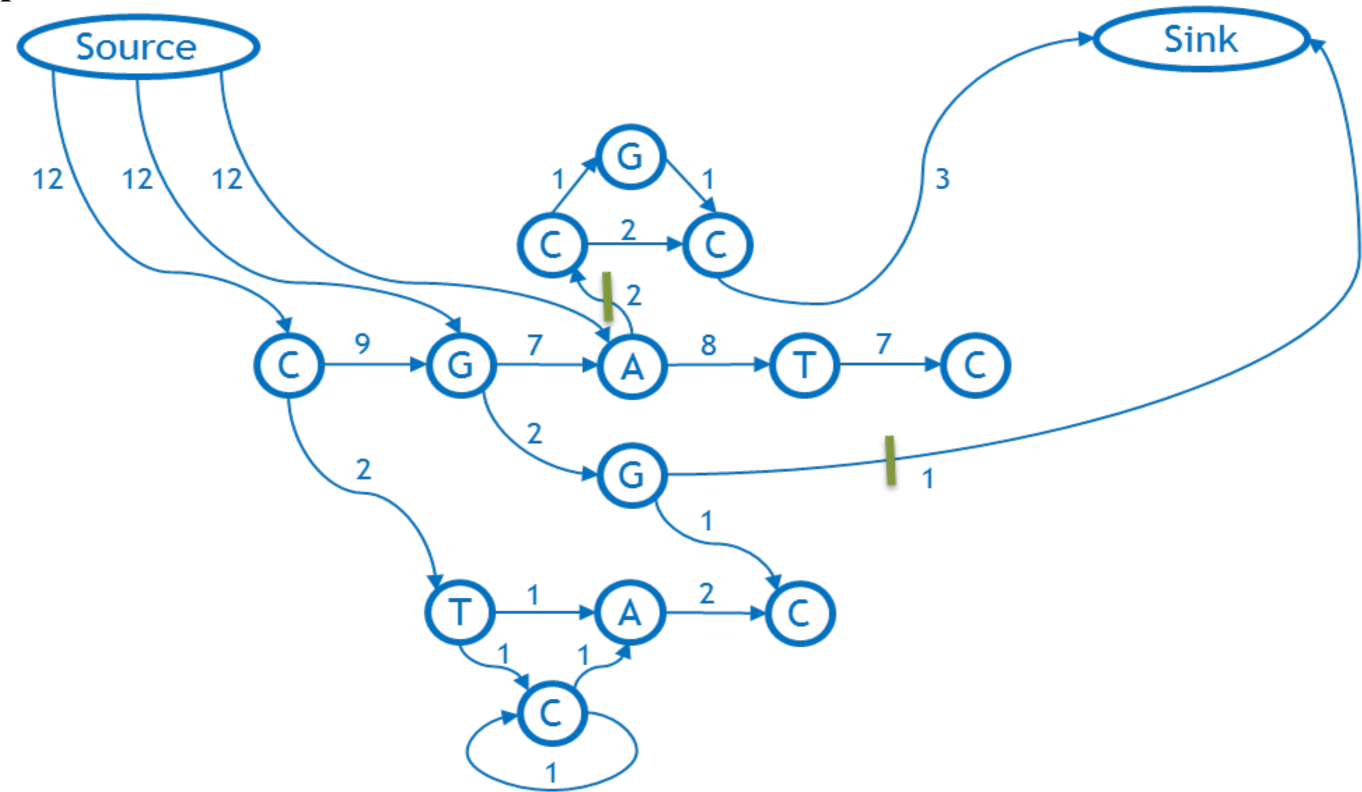
The flow network corresponding to the above weighted graph and its designated backbone.

We remove the cut arcs from the graph. Essentially, in doing this we decide to give up worrying about these arcs and try to minimize the weighted feedback and average cut width as if they were not there. Since capacity equals weight, by choosing a minimum capacity cut, we ignore the arcs that cost us the least in weight.

Excluding any initial feedback arcs on the backbone itself, we classify all bases not on the backbone into a sequence of **out-growths** and **in-growths** as follows. Starting at the last base on the backbone that has an out-arc, we define its out-growth to be all bases reachable by a forward directed path from that base. Then we move backwards along the backbone, defining out-growths consisting of all the bases reachable by a forward directed path from the base on the backbone with an out-arc that were not already included in previously defined out-growths. We define in-growths in a similar fashion, moving forward from the start of the backbone and using backward-directed paths. Figure 8 shows in-growth and out-growths for the example under consideration. There are two outgrowths, the first from the node G on the backbone (contains nodes G, G, C), and the second from its predecessor, the first node on the backbone (labeled “C”, contains nodes C, T, C, A). The dotted green arc is not a proper part of either outgrowth; it is discovered when exploring the outgrowth from C, and found to lead into the previous outgrowth found from G. The one in-growth enters the node A of backbone and contains nodes C, G, C, A, shown in purple.

**Figure 8.**
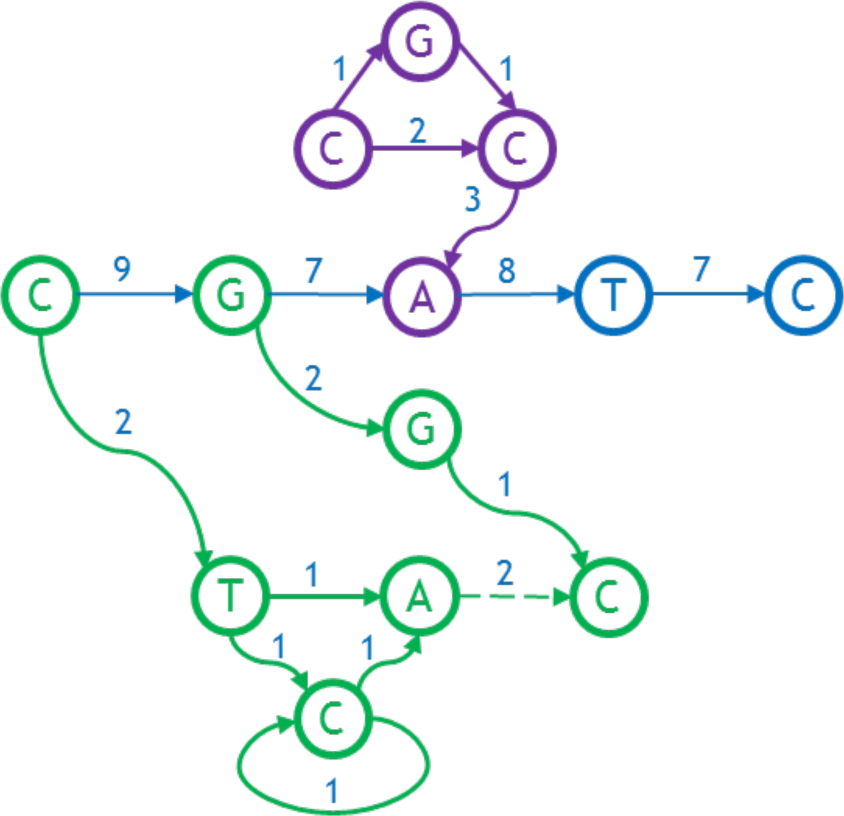
Out-growths (green) and in-growth (purple) of the graph.

### 4.4 Fourth Step. Repeat Procedure for In- and Out-growths

Finally, we apply the entire procedure recursively to each out-growth and in-growth, using a heuristic that uses the backbone’s base as the first base for an out-growth or as the last base for an in-growth, respectively. When the recursive call completes, the bases from it are inserted into the backbone of the calling procedure in the specified order immediately following an out-growth or immediately preceding an in-growth, respectively. The final ordering for the example above is shown in Figure 9.

**Figure 9.**
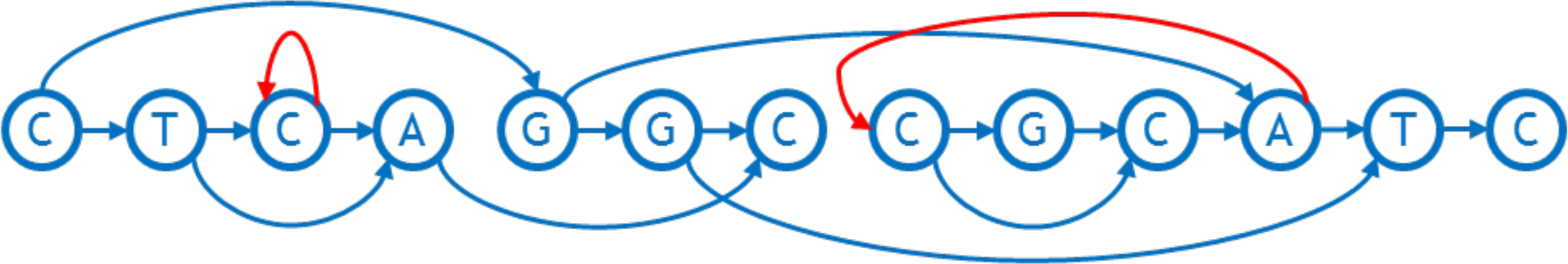
The final sorted graph with the bases totally ordered.

Normally the out-growths and in-growths together comprise the whole graph. However, if not, the entire procedure can just be repeated, each time using the linear order established from the previous cycle as a new backbone. Grooming a connected graph (see below) assures that these repetitions will eventually reach every node in the graph.

It remains to specify a heuristic for determining the backbone when it is not explicitly given. When the first base of the backbone is given, we extend it into a path using a greedy algorithm: in each step, we add to the existing path the base with the highest forward directed arc weight, breaking ties arbitrarily, and we do so until no more bases can be added. A complementary procedure is run in reverse if we are instead given the last base of the backbone.

### 4.5 Grooming

Finally, we explain the preprocessing step of grooming the graph. The base in each node in a sequence graph has two sides (3’ and 5’). The directed arcs we have been using are edges of the sequence graph that connect the 3’ side of one node to the 5’ side of another (possibly the same) node. The arcs are directed in the 3’ to 5’ side direction. There are also additional edges in a sequence graph that here we will refer to as **reversing joins**, which we have not discussed up until this point (see [1] and [9] for an introduction). A reversing join is an undirected edge connecting the 3’ sides of two nodes, or connecting the 5’ sides of two nodes. As a preprocessing step to the flow procedure, and all other heuristic algorithms we examine for linearization of a directed graph, we first eliminate as many of the reversing joins as possible by replacing the graph with an equivalent graph that has fewer reversing joins. Then if there are still reversing joins left we just ignore them. This way we are always working with graphs that only contain directed arcs. The process we use to minimize the number of reversing joins is called **grooming** (Figure 10).

**Figure 10.**
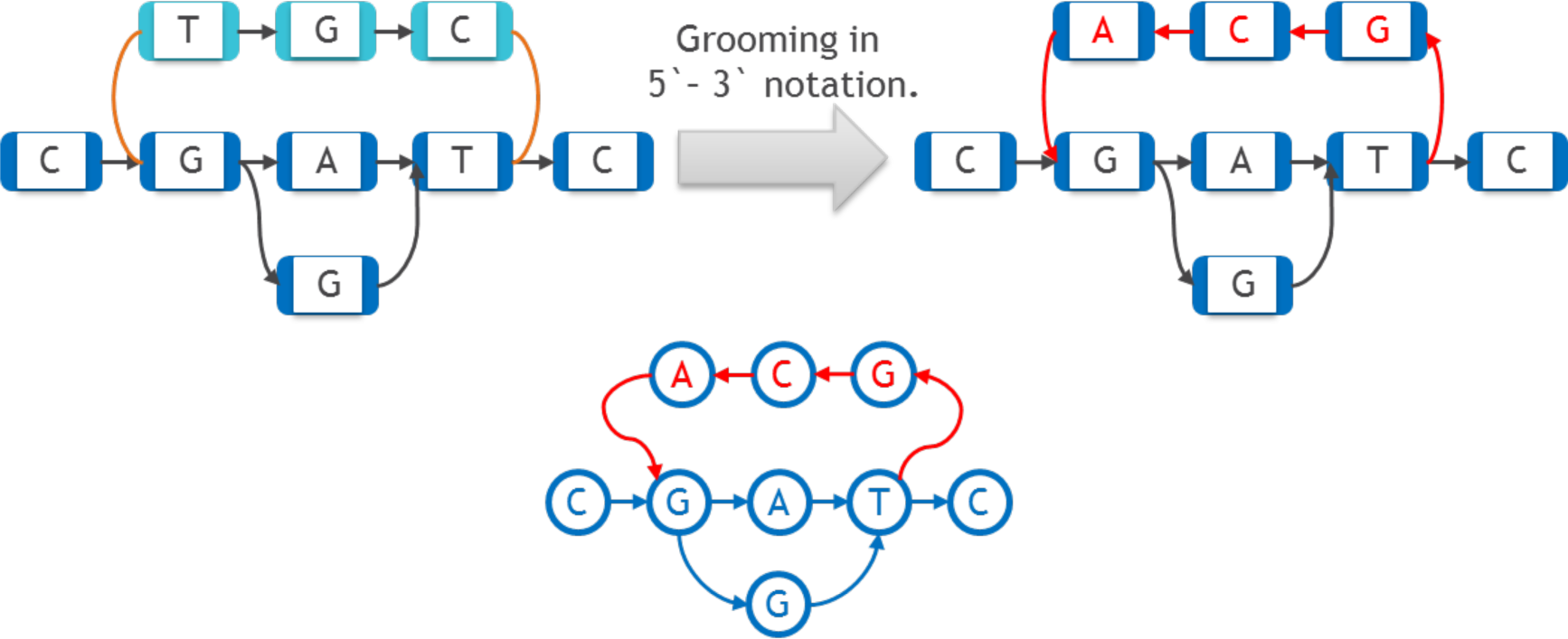
Grooming procedure. Reversing joins are shown in brown on the upper left. Bottom one shows the graph on the upper right as directed graph.

Grooming works as follows. A given connected component (a set of nodes such that one can travel between any two nodes in it along the standard arcs, in both direction, and reversing joins) may fall apart if the reversing joins are removed. This indicates that some of reversing joins were unnecessary. Let one connected component be called the **primary component** (shown in dark blue in Figure 10), and the others be called the **secondary components** (shown in light blue in Figure 10). We obtain an isomorphic graph that will have fewer components after removal of reversing joins by simply **reverse-complementing** the secondary component, i.e., reverse-complementing every base in it and inverting the direction of the arcs between these secondary bases. The 3’ reversing joins connecting this secondary component to the main graph are replaced by directed arcs pointing into the secondary component, and the 5’ reversing joins are replaced by directed arcs back to the primary component. This has the effect of changing every 3’ side in the secondary component into a 5’ side, and vice versa. On the right side of Figure 10 nodes and arcs which were changed during grooming are shown in red. By repeating this procedure on any connected sequence graph we eventually reach an isomorphic graph that has just one connected component even after removing the reversing joins.

Since we will not be reducing the number of reversing joins further, and being undirected they cannot be considered feedback arcs, from here on, we will stop paying attention to the reversing joins and consider only graphs that are fully directed.

## 5 Complexity Estimation

The max flow/min cut sorting algorithm described above can be broken into four steps:

1. Find the backbone (if it is not given)
2. Create the flow graph by adding the source and sink and connecting them to the graph
3. Find the maximum flow and minimal cut and delete the minimal cut, using the Ford-Fulkerson algorithm
4. Find the in-growth and out-growth and repeat steps 1 through 4 for them

Let us consider the complexity of each step separately.

In the preparation step, we perform a greedy depth-first search to find the backbone if it is not given (in practice we only find the backbone on recursive calls, as the whole graph’s backbone *is* given). We do not visit any arc more than once, so the time complexity is O(|A|), where A is the number of arcs.

In creating the flow, we add 2 nodes to the graph (the source and sink) and draw several arcs from the source to those nodes in the backbone that have an outgoing arc, and also reroute the arcs going into the backbone to the sink. We do not examine any arc more than once, so the time complexity is O(|A|).

The Ford-Fulkerson algorithm works in O(|A|*|max-flow|) [10]. In the worst case, |max-flow| ~ O(|A|), in which case the time complexity becomes O(|A|^2^).

Every recursive call decreases the number of nodes in the graph, so the number of recursive calls is O(|V|).

These estimations are shown in the table below.

The final complexity estimate is thus **O(|A|*|max-flow|) * |recursion depth|.**

The best-case complexity (if the max flow is constant and there are a constant number of recursive calls) is O(|A|), again, assuming at least one arc per node. In the worst case, it is O((|A|^2^) * |V|) as the max flow may be proportional to |A| and the recursion depth proportional to |V|.

The Ford-Fulkerson algorithm and the recursion contribute the most to the final algorithm’s complexity. Depth of recursion is not a problem in practice. However, the maximum flow will often be approximately O(|A|) due to how the flow is constructed: each variation increases it by creating a new path from the source to the sink. Because the flow becomes so large, the Ford-Fulkerson algorithm will work in quadratic time (O(|A|^2^)). Improvements to the algorithm that reduce the typical max flow so that it is polylogarithmic in |A| would improve its speed.

## 6 Experimental Evaluation

### 6.1 Data Modeling

The flow procedure was tested on data that was artificially generated by taking a 37 kilobase piece of the GRCh38 assembly and adding artificial structural variations to it using the RSVSim [11] from Bioconductor. This package lets one simulate any given set of structural variations to a reference, producing a modified FASTA file. The positions of the variations were distributed uniformly, while their lengths were fixed. After fixing a specified set of variations, a series of FASTA files were created and passed on to a vg [12] tool, which generated the graph using a multiple sequence graph alignment algorithm.

Four types of structural variation were simulated: insertions, deletions, duplications, and inversions. Tandem duplications were limited to one copy.

Two different test data sets, each consisting of a series of graphs, were generated in this way. The first was created to investigate the effect of the overall amount of variation on the number of feedback arcs and the cut width achievable by each algorithm. This data set consisted of graphs each having equal numbers of all four kinds of variations (i.e., there were as many insertions as there were deletions, inversions and duplications), with only the total number of variations changing between graphs. The variations’ sizes are given in Table 2.

**Table 2.** The lengths of the variations in the testing data.

|  | Deletion | Insertion | Inversion | Duplication |
| --- | --- | --- | --- | --- |
| Length | 20 | 20 | 200 | 500 |

The second set of graphs was created to investigate the relationship between the relative frequencies of each type of variation and the number of feedback arcs and the cut width achievable by each algorithm (Appendix B, tables 1–4).

**Table 1.**
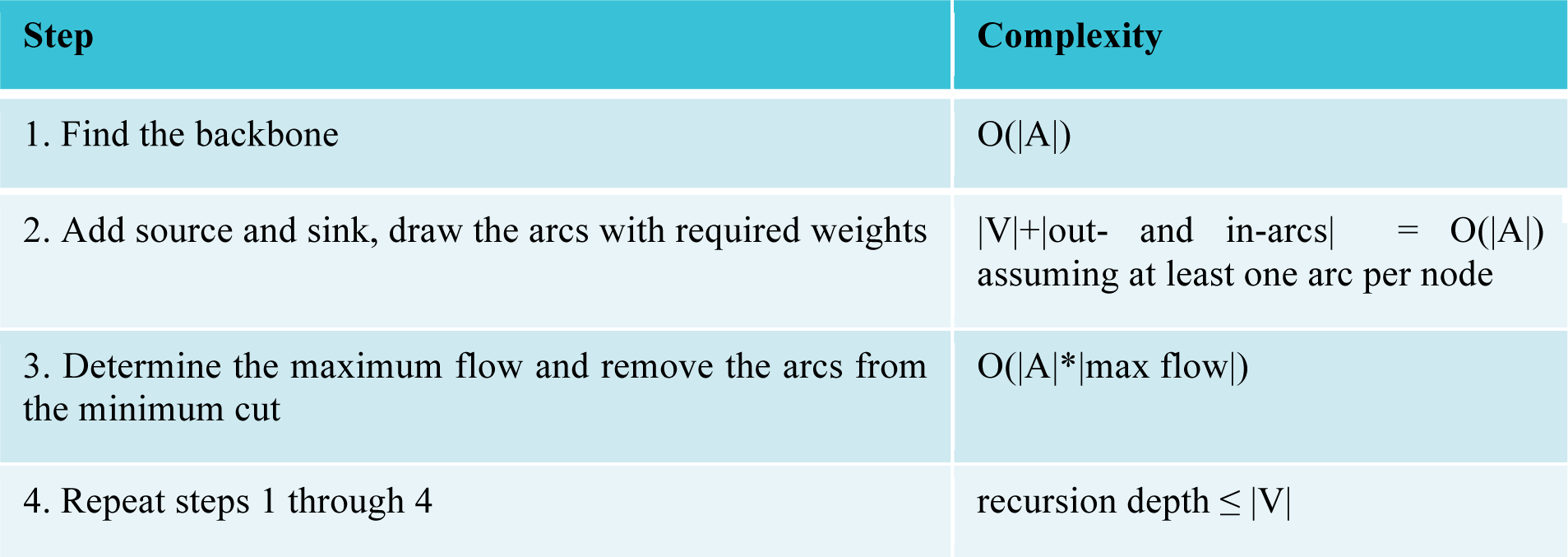
Complexity estimation of the flow procedure algorithm.

**Table 4.** The relationship between the number of feedback arcs and ACW, and the number of deletions.

| Number of Deletions | Kahn |  | Eades |  | Flow procedure |  |
| --- | --- | --- | --- | --- | --- | --- |
|  | Number of feedback arcs | Average cut width | Number of feedback arcs | Average cut width | Number of feedback arcs | Average cut width |
| 7 | 602 | 40.245 | 197 | 58.168 | 236 | 4.869 |
| 9 | 572 | 37.886 | 192 | 57.853 | 235 | 4.955 |
| 11 | 687 | 40.494 | 204 | 65.725 | 266 | 5.303 |
| 13 | 598 | 42.668 | 199 | 58.555 | 236 | 4.91 |
| 15 | 602 | 39.285 | 196 | 56.794 | 235 | 4.939 |
| 17 | 666 | 42.609 | 204 | 62.68 | 259 | 5.021 |
| 19 | 648 | 42.628 | 197 | 60.079 | 243 | 5.239 |
| 21 | 642 | 43.129 | 199 | 61.346 | 246 | 5.085 |
| 23 | 609 | 37.592 | 196 | 61.243 | 235 | 5.122 |
| 25 | 682 | 39.222 | 202 | 64.486 | 249 | 4.745 |
| 27 | 676 | 37.396 | 204 | 66.757 | 268 | 5.167 |
| 29 | 652 | 39.633 | 197 | 64.819 | 265 | 5.22 |
| 31 | 760 | 41.44 | 205 | 67.13 | 249 | 5.305 |

The third data set was created in order to test algorithm’s time performance. The graphs in this data set were created from scratch and have structure similar to those graphs above with doubling of the number of nodes from one to another.

### 6.2 Results and Discussion

In order to comparatively analyze the quality and speed of our algorithm, we took Kahn’s well-known topological sorting algorithm [13], as well as Eades’ [14] modified version thereof that guarantees a low number of feedback arcs while still working in linear time. We were not able to find competitor for cut width minimization problem to measure against, because all the algorithms we have found are focused on exact solution of the problem, thus having high complexity (cubic and higher) and thus working only with graphs of 200 nodes or less. Note that neither Kahn’s nor Eades’ algorithm uses the backbone as a heuristic. All three algorithms were tested on the same data (see Appendix B and Section 5.1 for details on the modeling). Their outputs were compared on two metrics – the number of feedback arcs and the average cut width. Kahn’s algorithm, Eades’ algorithm, and flow procedure are all implemented in the vg tool [12].

Figures 11 and 12 depict the main quantitative outputs of the three algorithms, Kahn, Eades, and the flow procedure (FP), for the first set of testing data. The same results for the second set are given in Appendix B. From Figure 11 it is fairly obvious that in terms of the number of feedback arcs, the FP algorithm vastly outperforms Kahn’s, doing only slightly worse than Eades’ algorithm. The difference between FP and Eades is much smaller than between FP and Kahn. It is clear from Figure 12 that FP’s average cut width is an order of magnitude lower than that of Eades [14] or Kahn [13].

**Figure 11.**
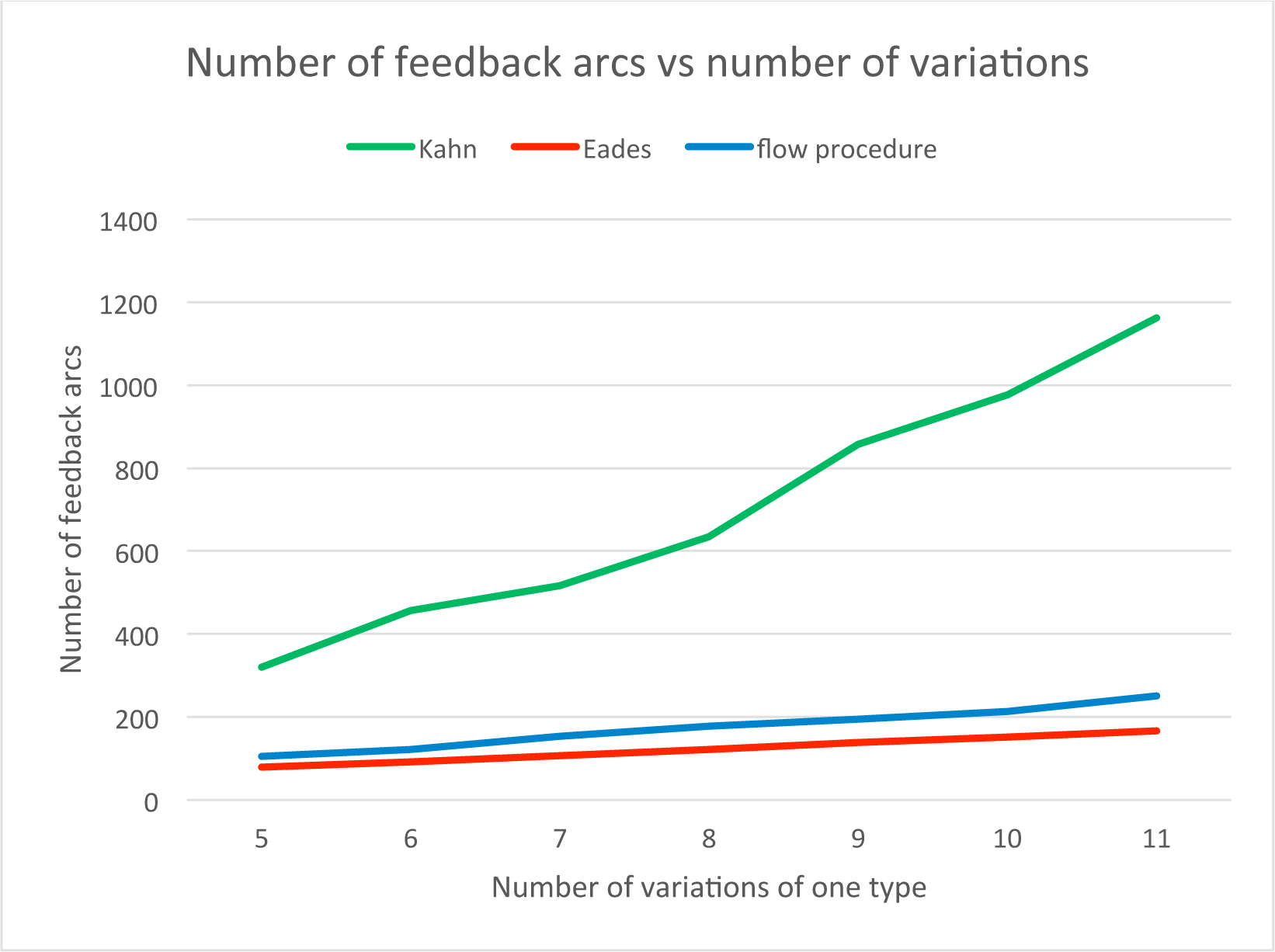
The relationship between the number of feedback arcs and variations

**Figure 12.**
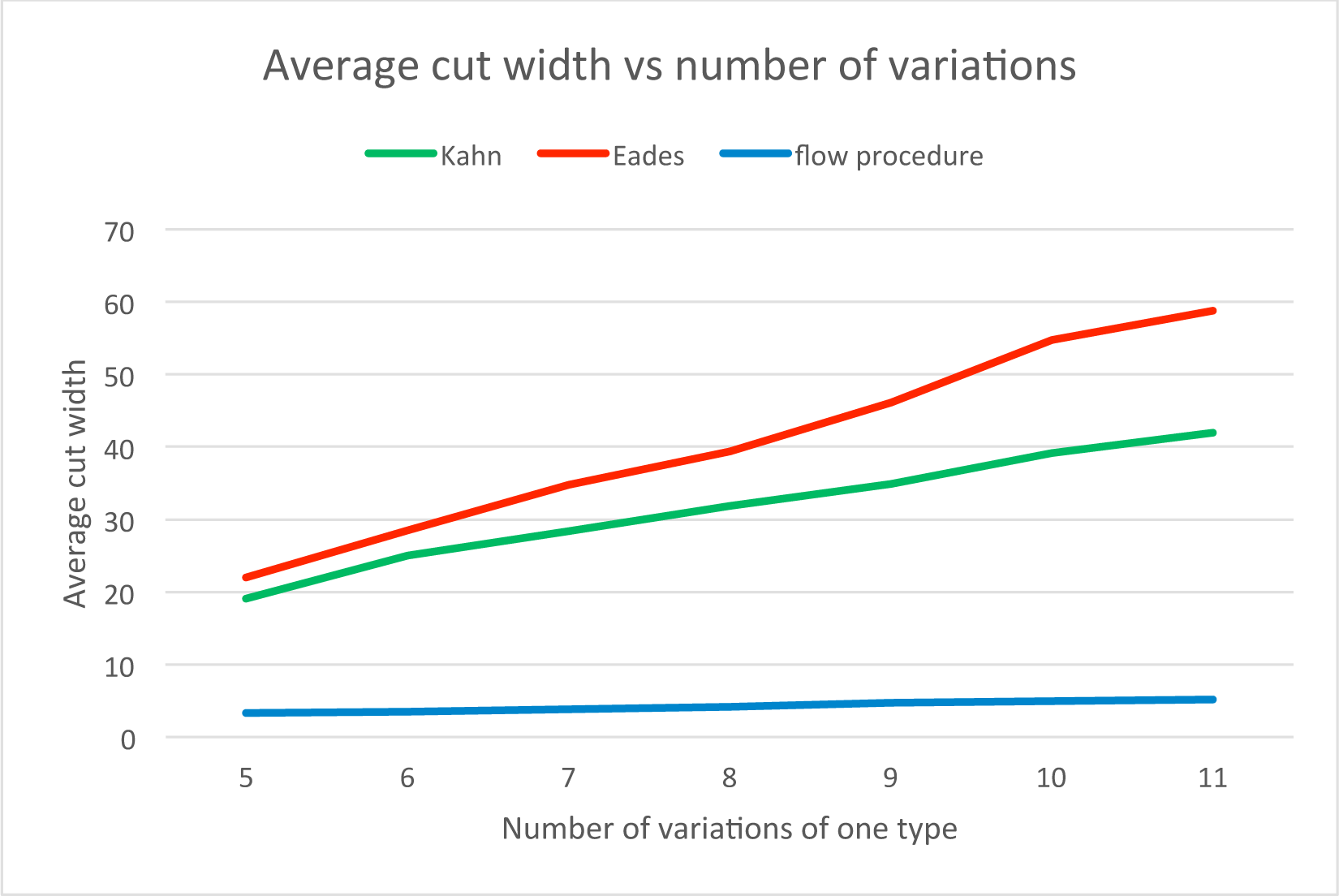
The relationship between the ACW and number of variations.

To use the FP algorithm in practice, one must estimate the time it takes the algorithm to run on large amounts of data. In order to use the algorithm on large graphs in practice, we would split the graph into pieces using a graph decomposition scheme as described in [2]. In practice the time complexity of the FP is O(|A|^2^). On the other hand, both Kahn’s and Eades’ algorithms have complexity O(|A|+|V|), since they do not pass any node more than twice. The relationship between the number of nodes in the graph and the algorithms’ runtimes on our test data is shown in Figure 13.

**Figure 13.**
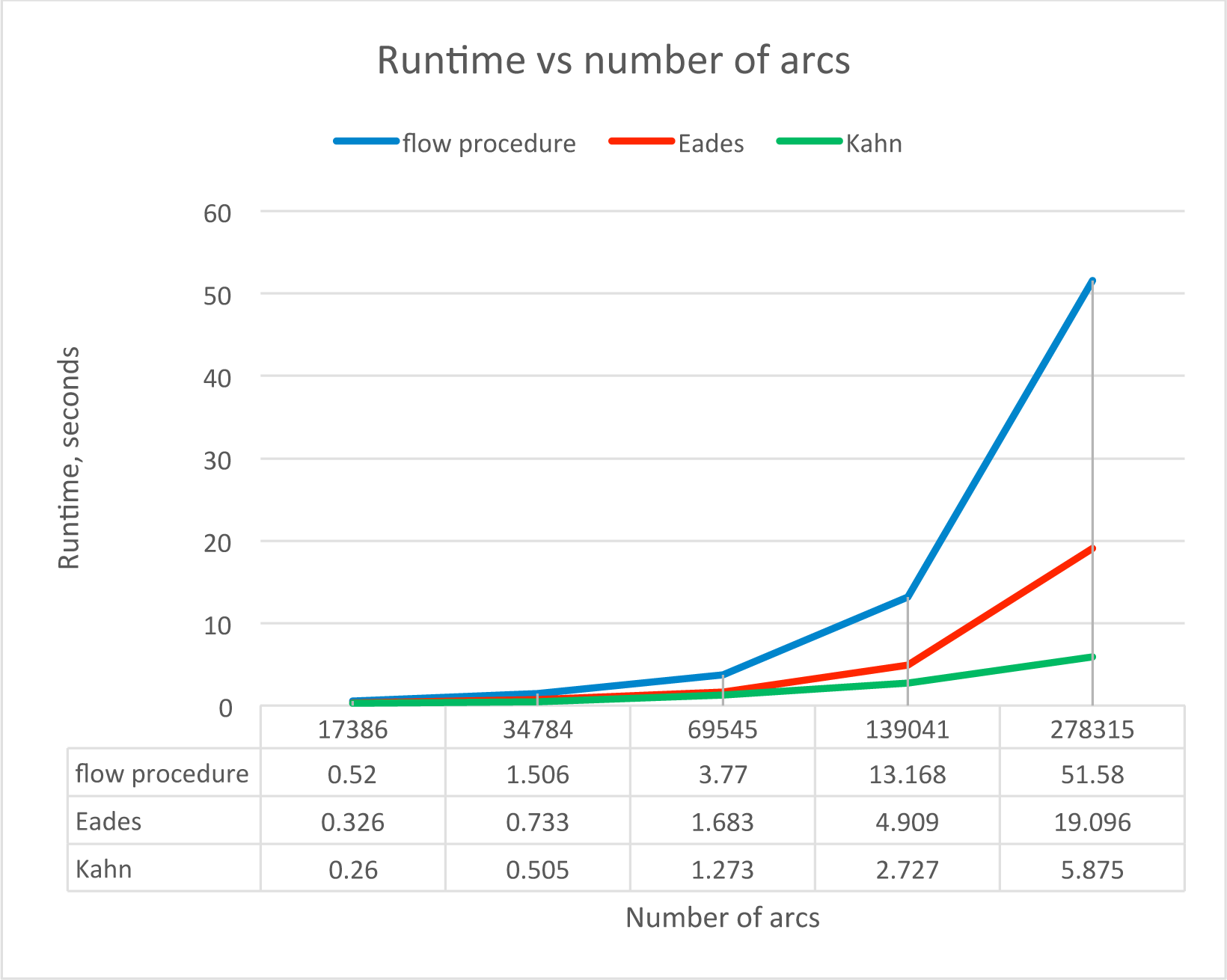
The relationship between the runtime and the number of arcs in the graph.

**Figure 14.**
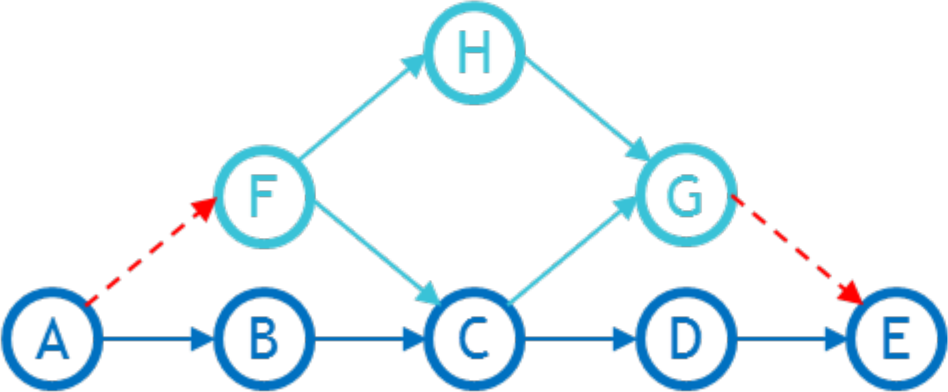
An example of a graph that requires a rerun.

**Figure 15.**
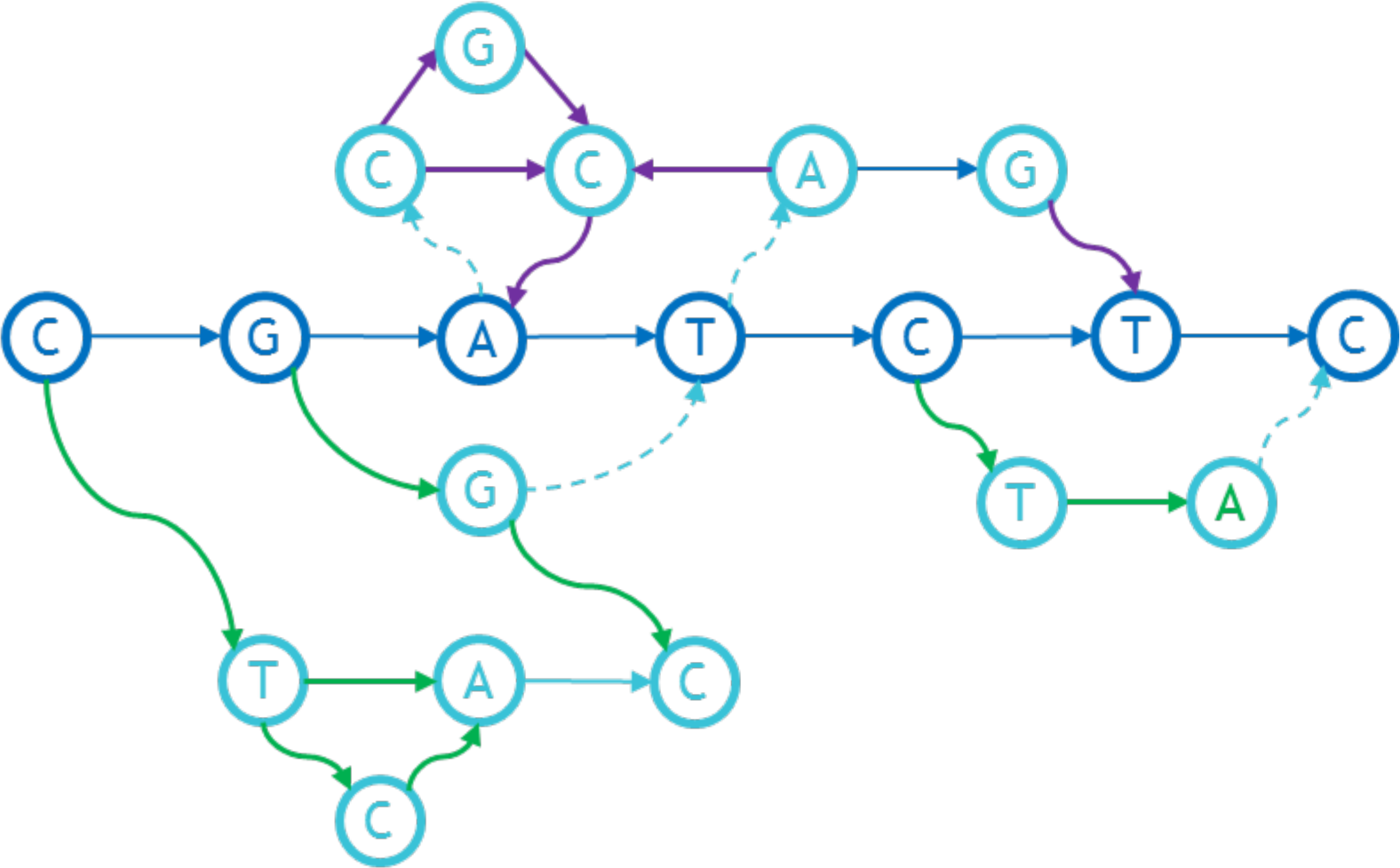
An example of the in-growths and out-growths for a graph.

**Figure 16.**
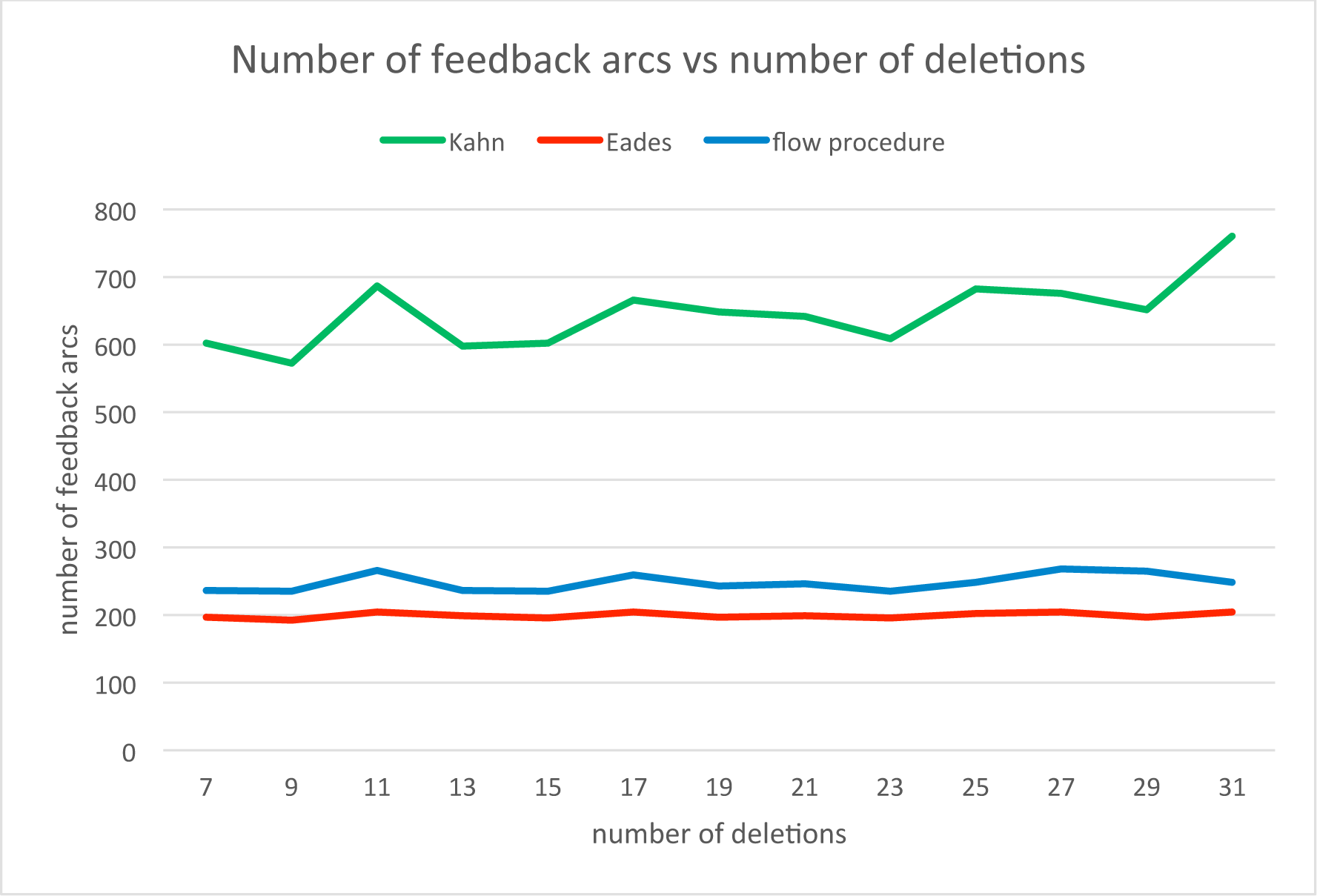
The relationship between the number of feedback arcs and the number of deletions.

**Figure 17.**
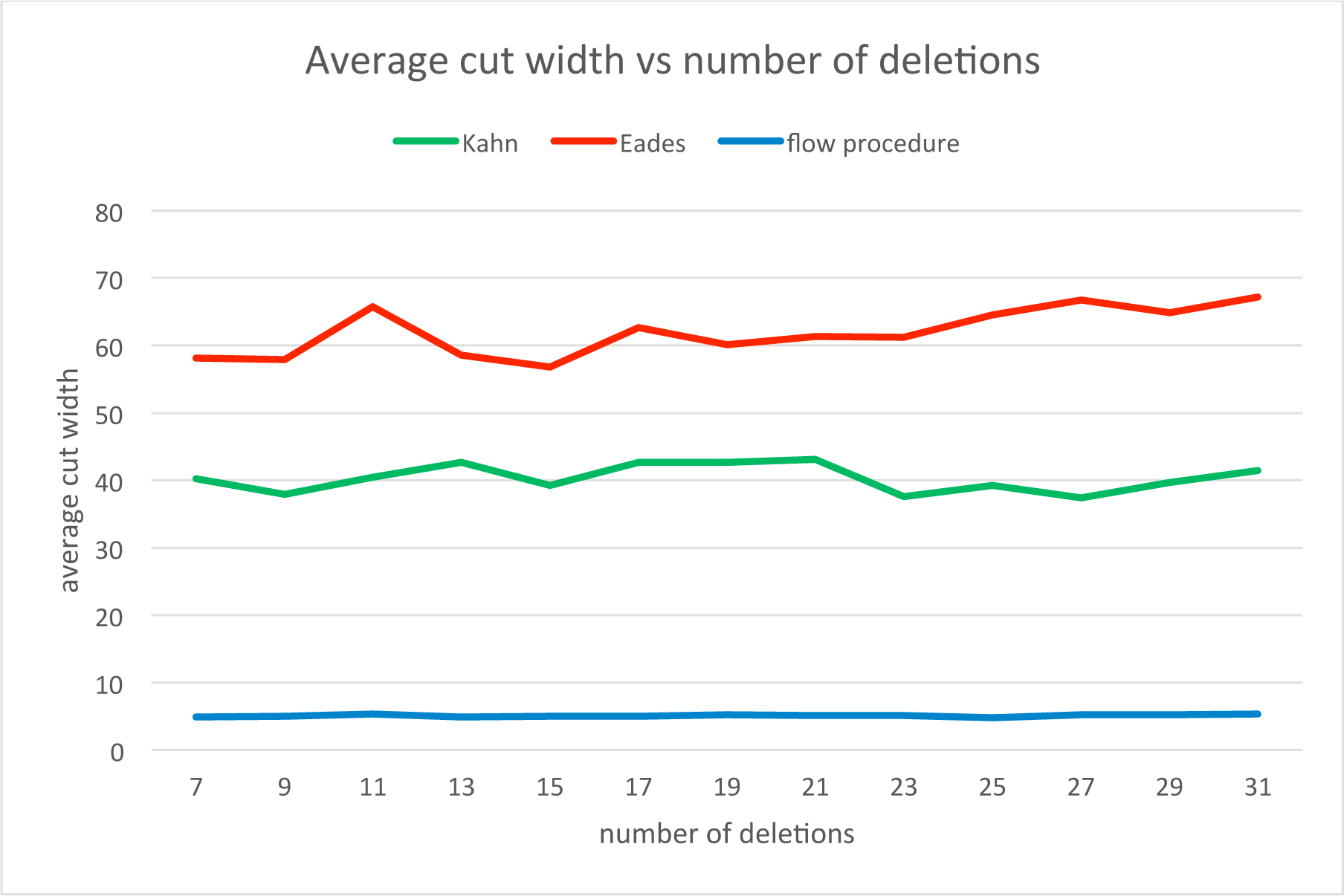
The relationship between the ACW and the number of deletions.

**Figure 18.**
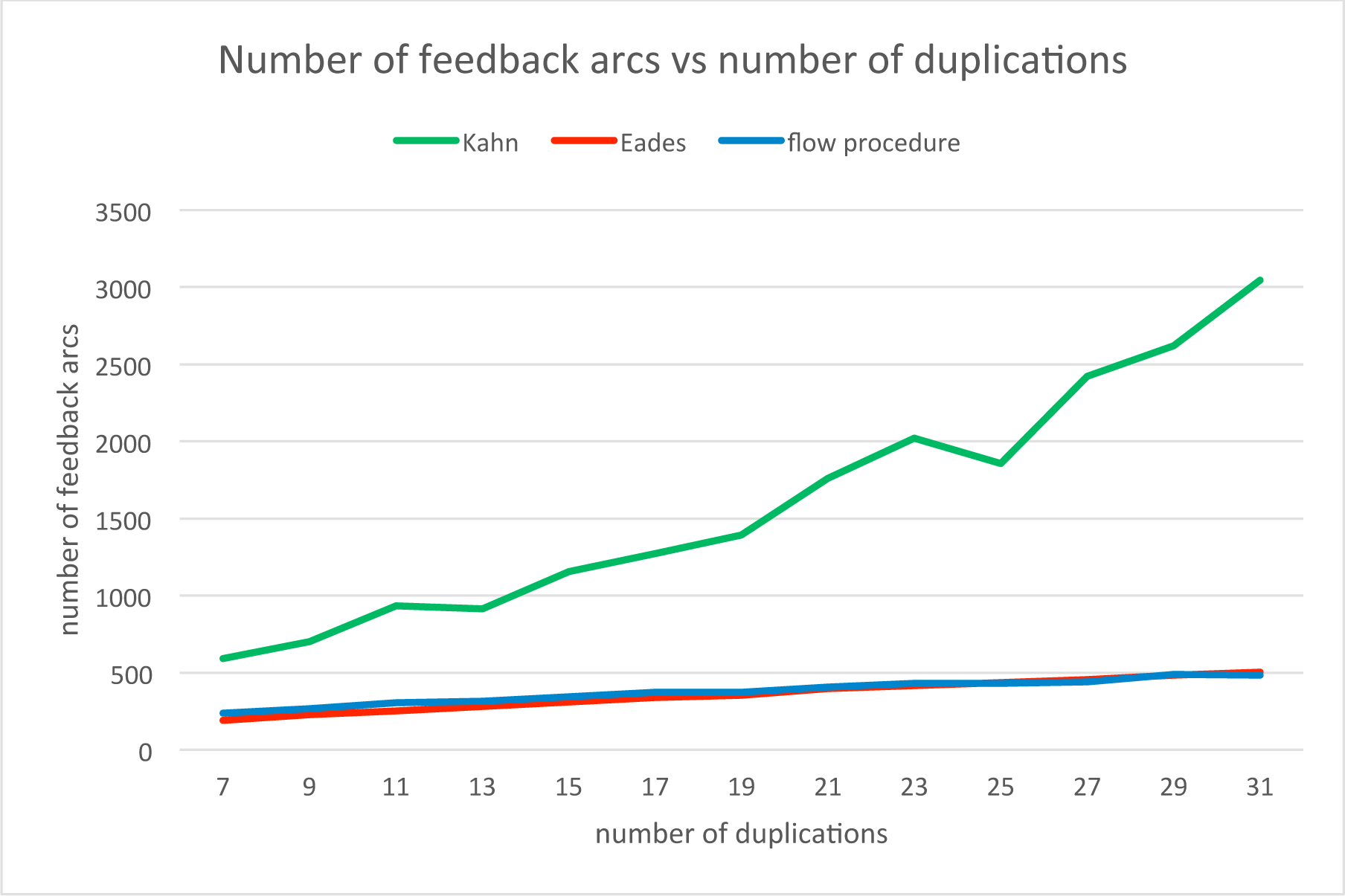
The relationship between the number of feedback arcs and the number of duplications.

**Figure 19.**
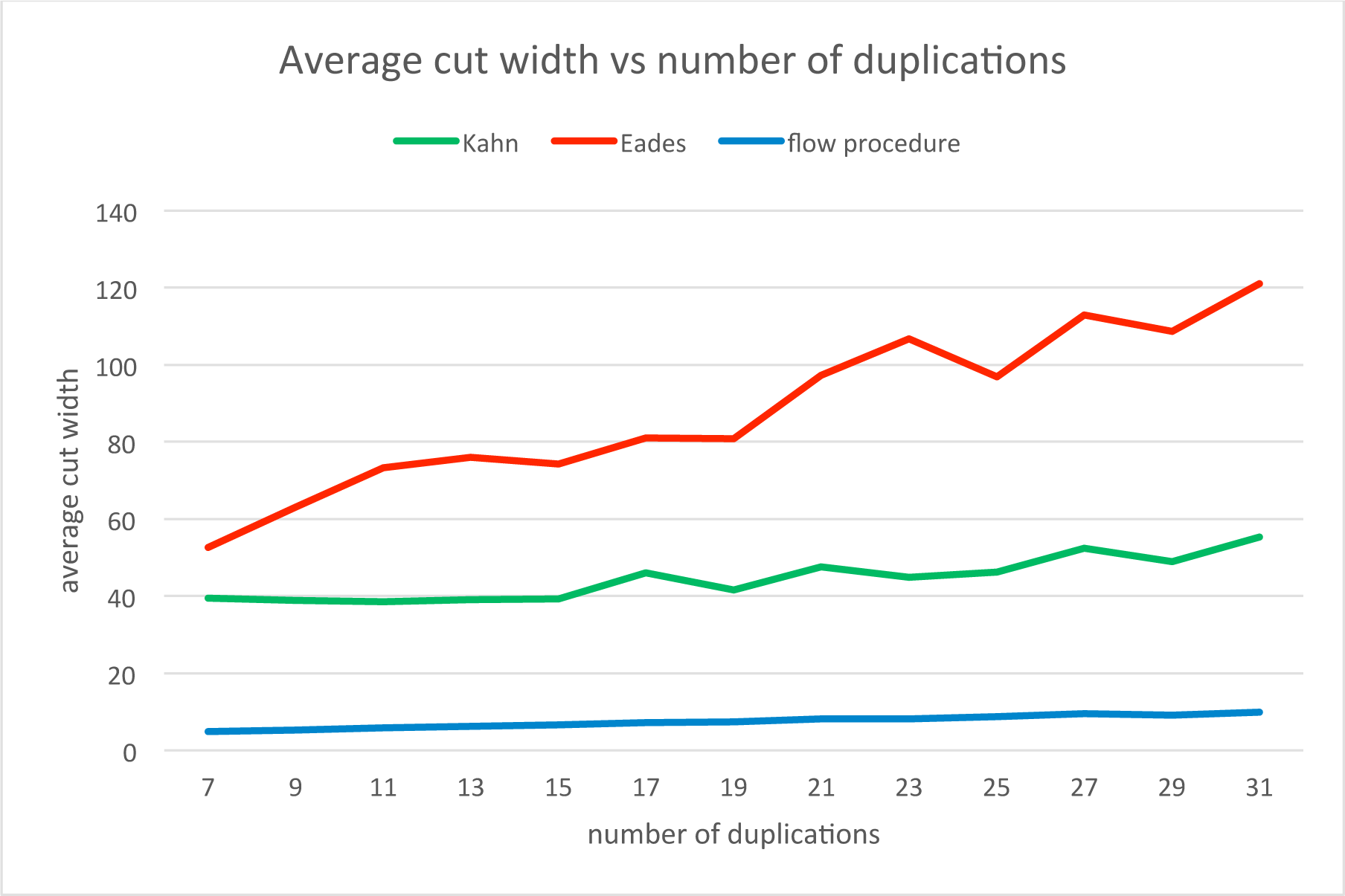
The relationship between the ACW and the number of duplications.

**Figure 20.**
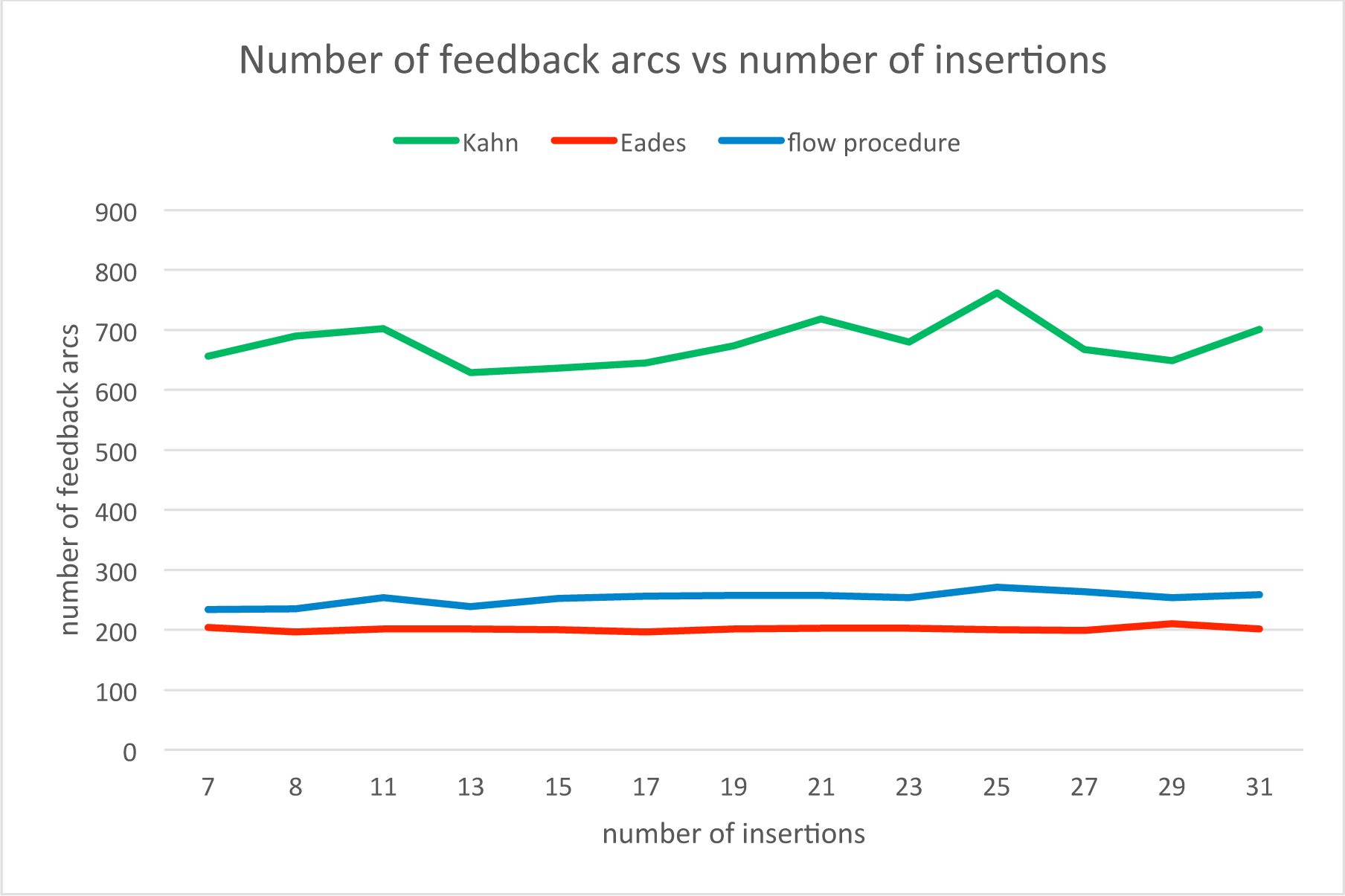
The relationship between the number of feedback arcs and the number of insertions.

**Figure 21.**
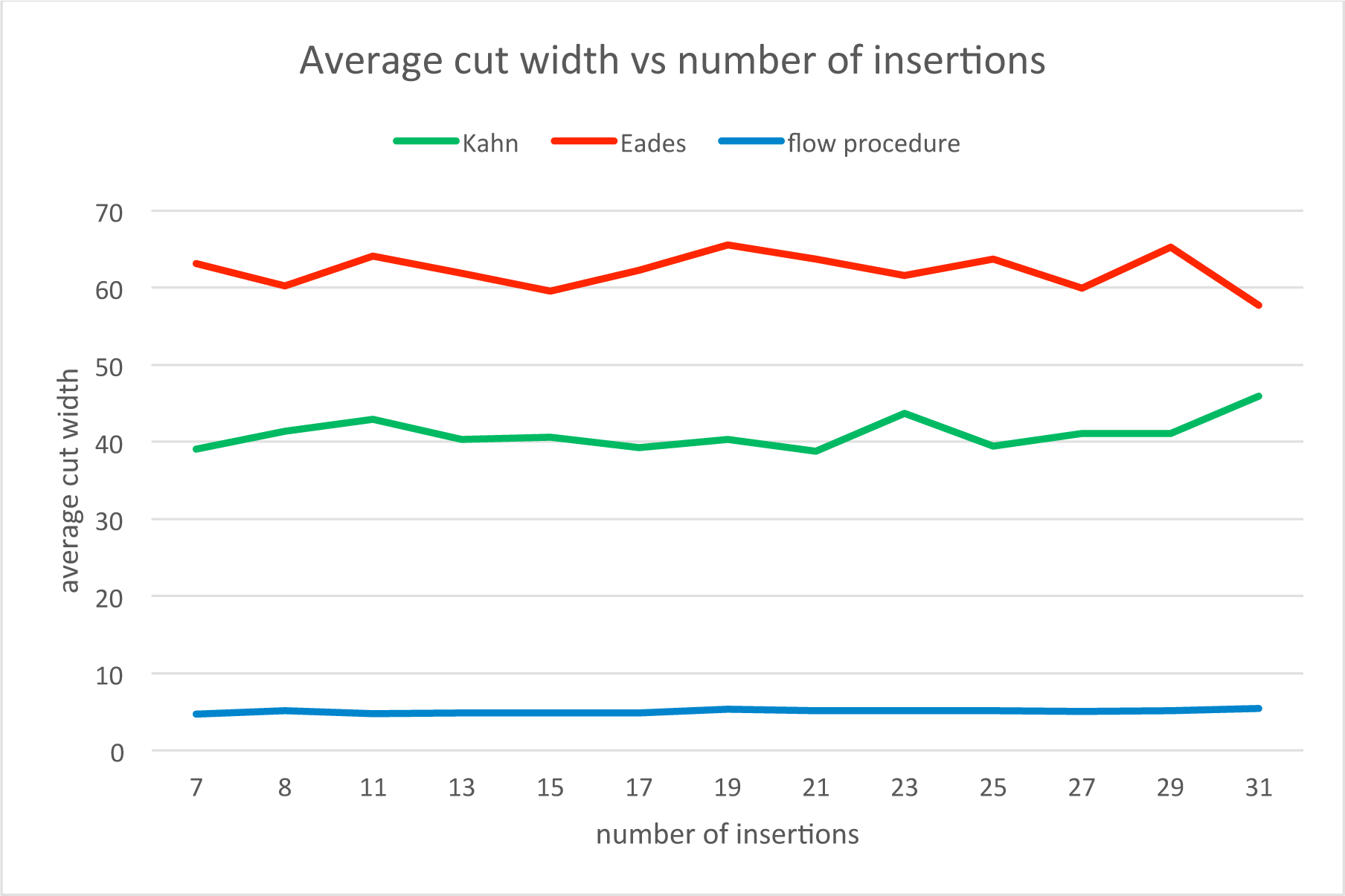
The relationship between the ACW and the number of insertions.

**Figure 22.**
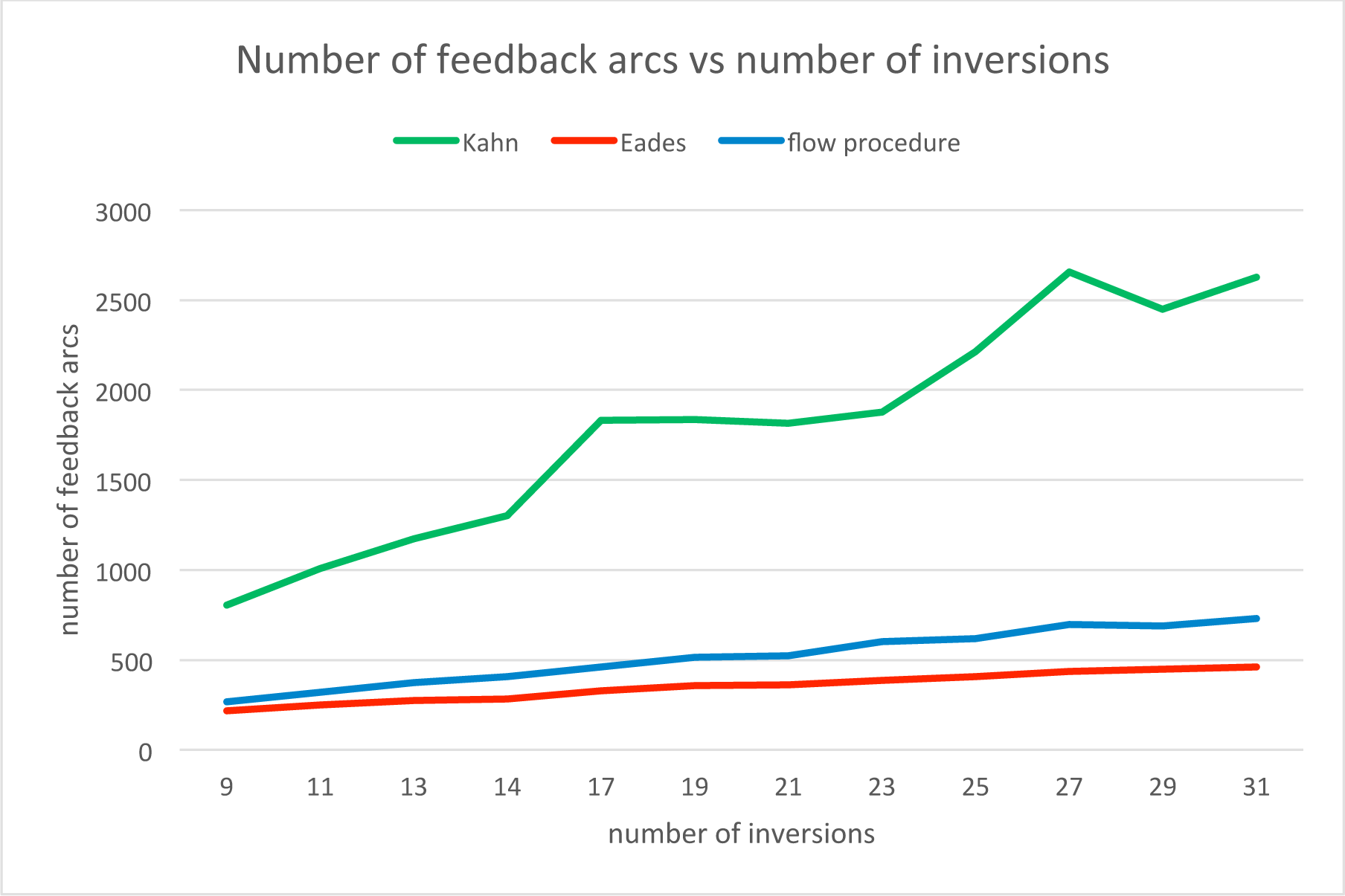
The relationship between the number of feedback arcs and the number of inversions.

**Figure 23.**
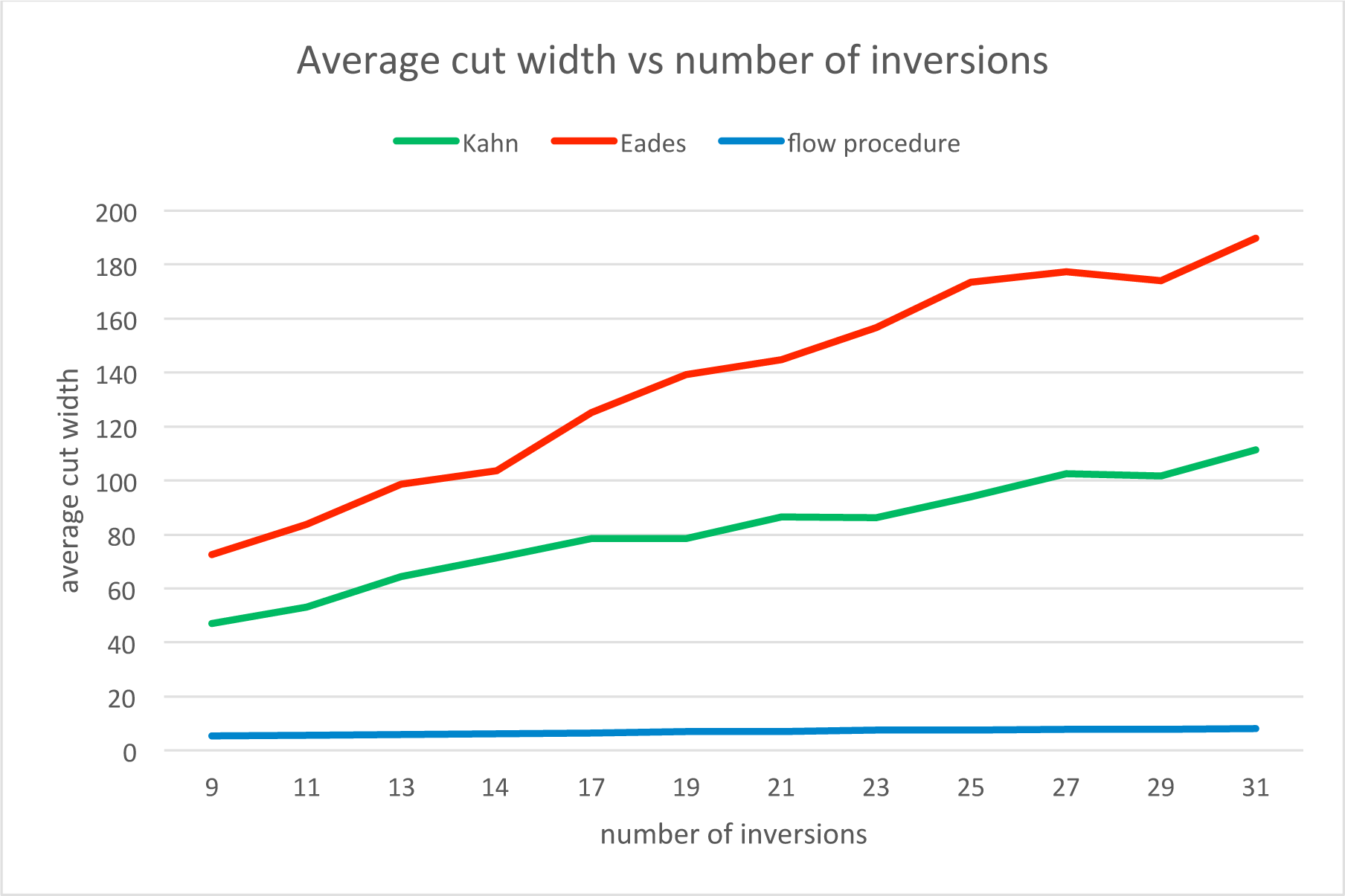
The relationship between the ACW and the number of inversions.

The relationship matches the one predicted theoretically. It shows that despite quadratic complexity estimation, it is clearly seen from Figure 13 the algorithm can be used on big graphs.

In addition to these tests on synthetic data, the algorithms were tested on a graph created from MHC region of chromosome 6 with 251297 nodes. Flow procedure running time was about 40 minutes.

## 7 Conclusion

We have proposed a new sequence graph linearization algorithm that outperforms standard methods on the criteria that are important for storing, traversing, analyzing and visualizing genome sequence graphs. The quantitative results thus obtained suggest that this algorithm will prove useful in genome exploration. Earlier work on sequence graph linearization [15] focused on minimizing feedback arcs, here we additionally introduce cut-width as an important measure of a linearization that effectively measures contiguity between elements that are connected. Future effort to lower the computational complexity of the algorithm using graph decomposition (see [2]) will allow us to apply a modified form of the presented algorithm to complete human scale sequence graphs of hundreds of millions of nodes.

## Acknowledgements

We’d like to thank Erik Garrison and Glenn Hickey for helpful conversations. This work was supported by the National Human Genome Research Institute of the National Institutes of Health under Award Number 5U54HG007990 and grants from the W.M. Keck foundation and the Simons Foundation. The content is solely the responsibility of the authors and does not necessarily represent the official views of the National Institutes of Health.

## 8 Appendix A

### 8.1 Some Details of the FP Algorithm

We pointed out in the algorithm description above that sometimes not all of the graph’s nodes end up in the final list, and so we need to rerun the procedure with the graph’s sorted part as the backbone. Fig. 1 demonstrates an example of such a situation.

Here *ABCDE* is the backbone (shown in dark blue), arcs AF and GE are in the min cut and hence deleted in the first run. *CG* is the in-growth and *FC* is the out-growth. Node *H* is in neither the in-growth nor the out-growth and so does not end up in the list on the first run of the procedure. On the rerun, however, the backbone will be *ABFCGDE* and so *H* will fall into F’s outgrowth.

Also noteworthy is the order in which we find the in- and outgrowths. First, we traverse the backbone from end to start, finding the outgrowth for each node, *then* we traverse it from start to end, finding the ingrowth. We include in the in- and outgrowths only those nodes that did not end up in any of the previous out- or ingrowths (see Fig. 2).

### 8.2 Step-by-step Algorithm Run

Let's start from the moment when we have already found and removed the minimum cut. We go from the beginning to the end over the backbone (CGATC) and find the in-growth CCGA (upper 3 nodes and A from the backbone). For this in-growth we run the entire flow procedure recursively. Looking for the backbone, we start from A and search for incoming max weight arcs. We get CCA, then run the min cut search and remove the CG arc. Then we recursively go to the CCA backbone from the beginning to the end; we are looking for the in-growth. We find GC. For it we run the procedure, which arranges these two nodes in the obvious way. We insert the result into the backbone CCA with the G before the second C (the one that had the in-arc). Thus, we get CGCA. All nodes of this part are sorted, so the recursion is finished and we insert the resulting in-growth into the backbone of the source graph. Inserting to the backbone we get CGCGCATC. There are no other in-growths, so we turn to search for out-growths. We go from the end to the beginning. We find the GGC out-growth. It includes 3 consecutive nodes, so the recursive procedure for it throws out a natural GGC order. We insert to the backbone and get CGGCCGCATC. Then we look for the next out-growth. We find the CTCA starting from the first node of the backbone. For it, we run the procedure recursively. It finds the backbone CTA, then removes the min cut, finds the in-growth CA and inserts its C before the A: CTCA. There are no other in- or out-growths, so this part of the algorithm is finished and we insert nodes to the original backbone, finally getting CTCAGGCCGCATC.

### 8.3 Test Data Set Modelling

In order to simulate the test data, we used the RSVSim package (version 1.14.0) from the Bioconductor software (Release 3.4). As a reference genome, we took BSgenome.Hsapiens.UCSC.hg38 (version 1.4.1), alternative branch chr13_KI270842v1_alt, which is 37287 nucleotides long. Using the simulateSV command of the RSVSim package, we modeled genome fragments of 10 individuals with a given set of variations. Resulting FASTA files were submitted to the entry of the msga command of the vg utility (https://github.com/vgteam/vg). As a result, we got a sequence graph (*.gfa format). This graph is an input to the commands vg sort-f (Eades) and vg sort (Flow procedure) of the vg utility (https://github.com/vgteam/vg). Finally, we got text files with graph nodes ordered by linearization using the Kahn, Eades, and flow procedure algorithms respectively. To analyze the algorithm, we created the original software to get the number of feedback arcs and the cut width in abovementioned sorts. To reduce the impact of accidents, we repeated the procedure 20 times for each set of variations and average the results.

We created variation sets as follows. In the modelled genome fragments, we added 5 variation types: insertions, deletions, duplications, inversions, and translocations. The positions of all variations were uniformly distributed over the simulation section of the genome. Twenty percent of the insertions were duplicating sections of the DNA. Translocations were modelled using the shoulder exchange mechanism. The lengths of insertions and deletions were 20 nucleotides; the length of inversion was 200 nucleotides; the length of duplications was 500. The number of variations of each type was equal to 5 in the first set, 6 in the second, 7 in the third, and so on up to 11 in the latest set of variations. The appendix provides a dependence of the number of feedback arcs and cut widths of number of variations of the same type. For this study, the number of variations of all types, except the examined, were fixed at level 7, and the number of investigated variations were changing according to the following list: 7, 9, 11, 13, 15, 17, 19, 21, 23, 25, 27, 29, and 31.

## 9 Appendix B

### 9.1 The Results for MHC Graph

**Table 3.** Number of feedback arcs and average cut width by all three algorithms for MHC graph.

| Kahn |  | Eades |  | Flow procedure |  |
| --- | --- | --- | --- | --- | --- |
| Number of feedback arcs | Average cut width | Number of feedback arcs | Average cut width | Number of feedback arcs | Average cut width |
| 1275 | 66.767 | 108 | 48.942 | 47 | 21.574 |

### 9.2 The Results of Additional Experiments

**Table 5.** The relationship between the number of feedback arcs and ACW, and the number of duplications.

| Number of | Kahn | Eades | Flow procedure |
| --- | --- | --- | --- |

| Duplications | Number of feedback arcs | Average cut width | Number of feedback arcs | Average cut width | Number of feedback arcs | Average cut width |
| --- | --- | --- | --- | --- | --- | --- |
| 7 | 592 | 39.392 | 191 | 52.631 | 237 | 4.848 |
| 9 | 701 | 38.977 | 228 | 63.121 | 267 | 5.307 |
| 11 | 936 | 38.551 | 252 | 73.24 | 306 | 5.813 |
| 13 | 913 | 39.167 | 280 | 76.01 | 314 | 6.213 |
| 15 | 1157 | 39.224 | 309 | 74.247 | 344 | 6.633 |
| 17 | 1273 | 46.042 | 341 | 81.009 | 371 | 7.286 |
| 19 | 1395 | 41.642 | 354 | 80.756 | 371 | 7.365 |
| 21 | 1760 | 47.581 | 396 | 97.255 | 405 | 8.088 |
| 23 | 2019 | 44.938 | 415 | 106.833 | 431 | 8.173 |
| 25 | 1857 | 46.156 | 435 | 96.822 | 433 | 8.738 |
| 27 | 2421 | 52.409 | 455 | 112.936 | 443 | 9.544 |
| 29 | 2622 | 48.912 | 482 | 108.725 | 491 | 9.036 |
| 31 | 3046 | 55.375 | 503 | 120.981 | 484 | 9.871 |

**Table 6.** The relationship between the number of feedback arcs and ACW, and the number of inversions.

| Number of Insertions | Kahn |  | Eades |  | Flow procedure |  |
| --- | --- | --- | --- | --- | --- | --- |
|  | Number of feedback arcs | Average cut width | Number of feedback arcs | Average cut width | Number of feedback arcs | Average cut width |
| 7 | 656 | 39.084 | 204 | 63.148 | 234 | 4.699 |
| 8 | 690 | 41.387 | 196 | 60.224 | 235 | 5.113 |
| 11 | 702 | 42.901 | 201 | 64.116 | 254 | 4.757 |
| 13 | 629 | 40.335 | 202 | 61.833 | 239 | 4.824 |
| 15 | 637 | 40.603 | 200 | 59.54 | 252 | 4.858 |
| 17 | 645 | 39.292 | 197 | 62.266 | 256 | 4.858 |
| 19 | 674 | 40.31 | 202 | 65.503 | 258 | 5.342 |
| 21 | 718 | 38.793 | 203 | 63.688 | 257 | 5.107 |
| 23 | 680 | 43.724 | 203 | 61.591 | 254 | 5.105 |
| 25 | 762 | 39.496 | 200 | 63.727 | 271 | 5.153 |
| 27 | 668 | 41.089 | 199 | 59.935 | 264 | 5.05 |
| 29 | 649 | 41.091 | 210 | 65.3 | 254 | 5.19 |
| 31 | 701 | 45.925 | 202 | 57.727 | 259 | 5.396 |

**Table 7.** The relationship between the number of feedback arcs and ACW, and the number of insertions.

| Number of Inversions | Kahn |  | Eades |  | Flow procedure |  |
| --- | --- | --- | --- | --- | --- | --- |
|  | Number of feedback arcs | Average cut width | Number of feedback arcs | Average cut width | Number of feedback arcs | Average cut width |
| 9 | 804 | 46.938 | 217 | 72.532 | 267 | 5.352 |
| 11 | 1006 | 52.943 | 251 | 83.811 | 321 | 5.654 |
| 13 | 1174 | 64.487 | 275 | 98.653 | 374 | 5.88 |
| 14 | 1300 | 71.296 | 284 | 103.571 | 406 | 6.189 |
| 17 | 1830 | 78.467 | 329 | 125.099 | 459 | 6.461 |
| 19 | 1836 | 78.408 | 356 | 139.293 | 515 | 6.873 |
| 21 | 1815 | 86.534 | 363 | 144.815 | 521 | 6.935 |
| 23 | 1876 | 86.29 | 385 | 156.763 | 601 | 7.406 |
| 25 | 2211 | 93.899 | 408 | 173.52 | 619 | 7.581 |
| 27 | 2657 | 102.554 | 436 | 177.482 | 698 | 7.672 |
| 29 | 2449 | 101.576 | 449 | 173.951 | 690 | 7.787 |
| 31 | 2629 | 111.228 | 459 | 189.837 | 728 | 8.071 |

## References

1. Benedict Paten, Adam Novak, David Haussler, Mapping to a Reference Genome Structure, eprint arXiv:1404.5010.

2. Benedict Paten, Adam M Novak, Erik Garrison, Glenn Hickey.: Superbubbles, Ultrabubbles and Cacti. Proceedings of RECOMB 2017.

3. Ali Baharev, Herman Schichl, Arnold Neumaer, Tobias Achterberg.: An exact method for the minimum feedback arc set problem.

4. Richard M. Karp.: Reducibility among combinatorial problems. In R. E. Miller, J. W. Thatcher, and J. D. Bohlinger, editors, Complexity of Computer Computations, The IBM Research Symposia Series, pages 85–103. Springer US, 1972.

5. Brandenburg, F. Hanauer, K.: Sorting Heuristics for the Feedback Arc Set Problem. Technical Report. Number MIP-1104, (2011).

6. Gavril, F.: Some NP-complete problems on graphs. In Proceedings of the 11th conference on information Sciences and Systems. pp. 91–95 (1977).

7. Martí, R., Pantrigo, J., Duarte, A., Pardo, E.: Branch and bound for the cutwidth minimization problem. Computers & Operations Research. 40, 137–149 (2013). doi: 10.1016/j.cor.2012.05.016

8. Cormen, T., Leiserson, C., Rivest, R., Stein, C.: Introduction to algorithms. Mit Press, Cambridge (Inglaterra) (2009).

9. Medvedev, P. Brudno, M: Maximum Likelihood Genome Assembly. Journal of Computational Biology. 16, 1101–1116 (2009). doi: 10.1089/cmb.2009.0047

10. Flows in networks. Princeton University Press. (1962)

11. https://www.bioconductor.org/packages/release/bioc/html/RSVSim.html

12. https://github.com/vgteam/vg

13. Kahn, A.: Topological sorting of large networks. Communications of the ACM. 5, 558–562 (1962). doi: 10.1145/368996.369025

14. Eades, P., Lin, X., Smyth, W.: A fast and effective heuristic for the feedback arc set problem. Information Processing Letters. 47, 319–323 (1993). doi: 10.1016/0020-0190(93)90079-O

15. Nguyen, N., Hickey, G., Zerbino, D., Raney, B., Earl, D., Armstrong, J., Kent, W., Haussler, D., Paten, B.: Building a Pan-Genome Reference for a Population. Journal of Computational Biology. 22, 387–401 (2015). doi: 10.1089/cmb.2014.0146

